# Oxygen-permeable microwell plates for high-throughput spheroid culture

**DOI:** 10.64898/2026.10.05.756623

**Authors:** Jun Takahashi, Hiroyuki Kato, Heather N. Zook, Hsun Teresa Ku, Kozo Iiba, Shunya Mizuno, Katsuhiro Esashika, Hirotake Komatsu

**Author notes:** **Corresponding author:** Hirotake Komatsu, MD, PhD, Transplant Surgery, Department of Surgery, University of California, San Francisco 513 Parnassus Avenue, San Francisco, CA 94143, USA.

## Abstract

Oxygen (O_2_) transport is a major limitation in high-density cell culture, particularly as three-dimensional (3D) spheroids are increasingly used to better mimic in vivo tissue environments. Here, we developed a poly(4-methyl-1-pentene) (PMP)-based microwell platform, leveraging its lower molecular absorption and greater mechanical stability than the commonly used O_2_-permeable polydimethylsiloxane (PDMS), while retaining high O_2_ permeability. We measured O_2_ permeability in 50-µm-thick PMP membranes and found approximately 200-fold greater O_2_ permeability than the conventional 1-mm polystyrene culture-plate material. PMP membrane-based microwell plates with 350-µm-diameter wells maintained spatial separation of pre-formed hepatocyte spheroids while supplying O_2_ from the plate bottom, synergistically enhancing spheroid oxygenation. Compared with non-O_2_-permeable plates, the platform reduced localized hypoxia and hypoxia-associated cell injury while improving spheroid viability and growth. Collectively, PMP-based O_2_-permeable microwell plates provide a promising, scalable, high-throughput platform for spheroid culture with multiple applications in tissue engineering, drug screening and beyond.

## Introduction

Two-dimensional (2D) cell culture systems, including monolayers and suspension cultures in well plates, have long served as foundational platforms for biological research, drug screening, and mechanistic studies due to their simplicity, scalability, and compatibility with high-throughput workflows. These systems enable rapid assessment of cellular responses to chemical compounds, genetic perturbations, and environmental cues. However, conventional 2D cultures often fail to recapitulate key aspects of tissue architecture and cell-cell interactions observed in vivo.^1, 2^ Three-dimensional (3D) spheroid cultures have emerged as complementary models that better mimic physiological microenvironments by promoting cell aggregation, extracellular matrix deposition, and spatial organization. As a result, they are valuable for modeling tumor biology, organoid development, and regenerative processes.^3–10^

Reproducing physiologically relevant microenvironments is essential for generating meaningful experimental outcomes that can translate to clinical applications, particularly in drug screening and regenerative medicine research. Among the various factors that regulate cellular function, O_2_ represents a primary requirement for maintaining cell viability, metabolism, and tissue homeostasis.^11–16^ In native tissues, O_2_ is continuously supplied through vascular networks; however, in conventional in vitro culture systems, O_2_ delivery relies primarily on passive diffusion from the surrounding atmosphere. Importantly, incubator O_2_ levels may not reflect pericellular O_2_ because oxygen availability depends on cellular consumption, cell density, medium volume, and vessel geometry, and inadequate O_2_ delivery can alter metabolism and function across multiple cell types, including hepatocytes and cardiac organoids.^13, 17, 18^

Most culture platforms use non-O_2_-permeable polystyrene culture plates, which restrict direct O_2_ transport to cells. This limitation becomes increasingly pronounced in high-density cultures and 3D constructs such as spheroids, where cellular O_2_ consumption is substantially elevated. As a result, O_2_ consumption can exceed O_2_ delivery, leading to reduced pericellular O_2_ tension and the formation of hypoxic regions.^13, 17, 18^ Such O_2_ limitation can alter cellular metabolism and function, potentially reducing the physiological fidelity and translational relevance of in vitro culture models.^18^

Microwell platforms have been developed to generate spheroids from single cells and help to spatially organize pre-formed spheroids during culture, maintaining them in stable and controlled configurations. Such systems offer improved control over spheroid size and distribution and have demonstrated enhanced viability in 3D culture applications.^19–24^ However, most existing microwell platforms are also fabricated from non-O_2_-permeable plastics. While these platforms can regulate spheroid positioning and uniformity, they do not inherently address O_2_ transport limitations, leaving spheroids susceptible to diffusion-driven hypoxia.

To overcome these challenges, O_2_-permeable materials, particularly silicone elastomers such as polydimethylsiloxane (PDMS), have been introduced as alternative culture substrates to enable direct O_2_ diffusion from the bottom surface.^25–31^ PDMS-based platforms have demonstrated improved O_2_ availability and enhanced cellular function in various applications. However, PDMS exhibits substantial sorption of hydrophobic small molecules, which can alter effective molecular concentrations within the culture microenvironment and compromise experimental accuracy.^32–35^ These limitations highlight the need for robust, scalable O_2_-permeable culture materials that support precise O_2_ control without compromising compatibility with high-throughput biological assays. Among gas permeable materials, poly(4-methyl-1-pentene) (PMP) is a transparent and chemically stable polymer characterized by high O_2_ permeability and strong mechanical robustness.^36–41^ PMP has been widely utilized in industrial and medical fields, including gas separation membranes, medical device components, diagnostic systems, and optical materials, owing to its ability to permit efficient gas transport while maintaining liquid barrier properties.^42, 43^ In addition to its O_2_ permeability, PMP exhibits low sorption of small molecules and chemical compounds,^37, 44^ which addresses a known limitation of silicone-based materials, including PDMS. This property is particularly advantageous for studies involving hydrophobic or lipophilic compounds, and PMP membrane-based cell culture vessels have been successfully applied to nephrotoxicity assessments of lipophilic drugs.^44^ These features position PMP as an attractive material for scalable O_2_-permeable culture systems.

Accordingly, this study introduces PMP-based, O_2_-permeable 2D and 3D culture platforms supporting adherent, suspension, and spheroid systems, with broad applicability in drug screening, biological research, regenerative medicine, and tissue engineering.

## Materials and methods

### O_2_-permeable flat plates

Flat O_2_-permeable culture plates were obtained from Mitsui Chemicals (InnoCell^™^, Mitsui Chemicals, Inc., Tokyo, Japan; #FP0024C and FP0096C for collagen-coated surfaces, and #FP0024N and FP0096N for suspension culture). The culture plate bottom was composed of PMP, a material with high O_2_ permeability and low autofluorescence. The 50-µm-thick PMP membrane was fabricated using a melt-based film-forming process with controlled thickness and served as an O_2_-permeable bottom surface for cell culture.

### O_2_-permeable microwell plates

Microwell O_2_-permeable culture plates were fabricated by forming arrays of hemispherical microwells directly on the PMP membrane. Each microwell had a diameter of approximately 350 µm and a depth of 85 µm, while the membrane thickness at the base was comparable to those of the O_2_-permeable flat PMP membrane plates. Microwell structures were generated using a thermoforming-based process. To generate non-O_2_-permeable control conditions, a gas-impermeable polyethylene terephthalate (PET) film was adhered to the bottom surface of the PMP membrane, which prevents gas diffusion through the membrane. Overview images of the entire well area in a 24-well plate and cross-sectional images were acquired using an all-in-one fluorescence microscope (BZ-X710, KEYENCE CORPORATION, Osaka, Japan). 3D surface topography of the microwell structures was subsequently obtained using a digital microscope (VHX-X1, KEYENCE CORPORATION, Osaka, Japan) to characterize the architecture of individual microwells.

### O_2_ permeability measurement of culture plate materials

O_2_ permeability of the materials used in the culture plates was measured using a differential pressure gas permeability measurement system (BT-3, Toyo Seiki Seisaku-sho Ltd., Tokyo, Japan) at 23°C under 0% relative humidity, based on the general principles of ISO 15105-1:2007. The test sample was mounted between two chambers of the permeability measurement apparatus. One chamber was filled with O_2_ at atmospheric pressure, while the opposite chamber was evacuated to create a pressure gradient across the sample. O_2_ permeating through the material was detected by monitoring the pressure increase in the downstream chamber over time. O_2_ transmission rate was calculated from the steady-state gas flux. O_2_ permeability was then calculated according to the ISO standard and expressed as cm^3^/(m^2^·24 h·atm).

### K562 suspension cell culture

K562 cells (human chronic myelogenous leukemia cell line; provided by the RIKEN BRC through the National BioResource Project of the Ministry of Education, Culture, Sports, Science and Technology, Japan) were cultured in Roswell Park Memorial Institute (RPMI) 1640 medium supplemented with 10% fetal bovine serum, 1% GlutaMAX^™^ Supplement (Thermo Fisher Scientific Inc., MA, USA), 100 units/mL penicillin G and 100 µg/mL streptomycin sulfate (Penicillin-Streptomycin Solution (×100), FUJIFILM Wako Pure Chemical Corporation, Osaka, Japan), at 37°C in a humidified incubator with 5% carbon dioxide (CO_2_). Cells were seeded on flat O_2_-permeable plates at densities of 5 × 10^5^ cells/cm^2^ for medium O_2_ measurements under high density suspension culture conditions using a 24-well plate format (InnoCell^™^ #FP0024N), and 2 × 10^5^ cells/cm^2^ for cell proliferation assays using a 96-well plate format (InnoCell^™^ #FP0096N). Culture conditions included either daily partial medium exchange (one-third of the total volume) or no medium replacement throughout the 3-day culture period. Cell line passages 10-20 were used in all experiments.

### HepG2 monolayer culture

HepG2 cells (human hepatocellular carcinoma cell line; provided by the RIKEN BRC through the National BioResource Project of the Ministry of Education, Culture, Sports, Science and Technology, Japan) were cultured in Dulbecco’s Modified Eagle Medium supplemented with 10% fetal bovine serum, 100 units/mL penicillin G and 100 µg/mL streptomycin sulfate (FUJIFILM Wako Pure Chemical Corporation), at 37°C in a humidified incubator with 5% CO_2_. Cells were seeded on collagen-coated flat O_2_-permeable plates at densities of 4 × 10^4^ cells/cm^2^ for medium dissolved O_2_ measurements using a 24-well plate format (InnoCell^™^ #FP0024C), and onto 96-well plates (InnoCell^™^ #FP0096C) for cell proliferation assays in 2D monolayer culture experiments. Culture conditions included either daily partial medium exchange (one-third of the total volume) or no medium replacement throughout the 3-day culture period. For the medium O_2_ measurements on day 1, a no-cell control condition (medium-only wells) was also prepared. Cell line passages 10-20 were used in all experiments.

### HepG2 spheroid formation and culture conditions

HepG2 spheroids were generated using a conventional microwell platform (EZSPHERE 900SP, AGC Techno Glass Co., Ltd., Shizuoka, Japan). Single-cell suspensions were seeded at a density of approximately 5 × 10^5^ cells per well in a 6-well plate and cultured for 2 days to allow spheroid formation (approximately 150 µm in diameter). Using the HepG2 spheroids, two types of experiments were performed: spheroid culture on flat plates and culture on microwell plates. For both studies, formed spheroids were collected and transferred to either O_2_-permeable plates or non-O_2_-permeable plates with a gas barrier film attachment. For flat plate experiments, a 2 × 2 experimental design was employed, including spheroid density (200 or 600 spheroids per well in a 24-well format) and spheroid distribution (dispersed distribution versus central clustering within a 24-well plate). For microwell plate experiments, 600 spheroids were uniformly seeded within each well. The number of spheroids was manually counted prior to seeding. To visualize spheroid distribution within the microwell plate, spheroids were stained with Calcein acetoxymethyl ester (Calcein-AM) (2 µg/mL, DOJINDO Laboratories, Kumamoto, Japan). Fluorescence images covering the entire 24-well area were acquired using tile imaging (BZ-X710, KEYENCE).

### Dissolved O_2_ measurement in culture media

Dissolved O_2_ levels in the culture media were measured using a fluorescence-based optical O_2_ meter (FireSting-O2; FSO2-4) equipped with a robust O_2_ probe (PyroScience GmbH, Aachen, Germany). The O_2_ probe was held in place using a polytetrafluoroethylene (PTFE) tube adapter (F-5005-02, FLON INDUSTRY, Tokyo, Japan), and the measurement height was adjusted with a precision lab jack equipped with a digimatic indicator (LABZ-70-G, FINESYS, INC, Tokyo, Japan). The probe tip was directly inserted into the medium near the cell or spheroid layer. The entire measurement setup, including the O_2_ probe and probe holder, was placed inside a CO_2_ incubator maintained at 37°C. Continuous O_2_ tension was recorded immediately after attachment of the gas barrier film and subsequently for four hours. Data points were recorded every one minute during the measurement period. In addition, a separate measurement was obtained one day after culture to determine the equilibrated dissolved O_2_ level in the culture media. Data were collected using Pyro Oxygen Logger V3.2.1 software (PyroScience GmbH, Aachen, Germany) for subsequent analysis.

### Simulation of pO_2_ distribution

A finite element method (COMSOL Multiphysics 5.3, Stockholm, Sweden) was used to examine the O_2_ gradient formed in the microenvironment of the culture media in a 24-well plate. The O_2_ transport properties were characterized using Fick’s diffusion equation with the consumption term:

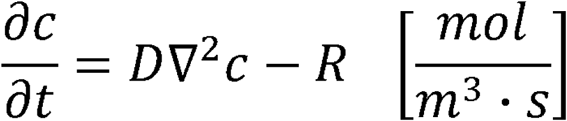

where *c* is the O_2_ concentration,*D* is the diffusion constant of O_2_, and *R* is the O_2_ consumption rate (OCR) of cells governed by the Michaelis-Menten type metabolic kinetics:

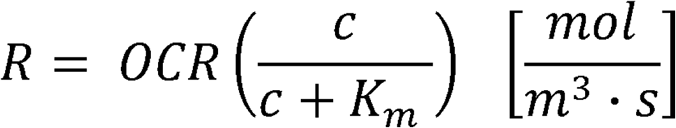

where *OCR* denotes the maximum OCR achievable by the spheroids and *K_m_* represents the O_2_ concentration at which the rate equals to half of the maximum. Henry’s law was also used to correlate the O_2_ concentration with O_2_ tension so the continuity of O_2_ partial pressure (pO_2_) can be applied across different boundaries:

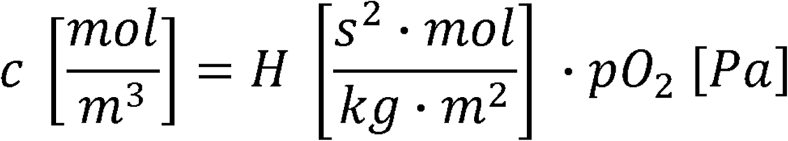

Where *H* is the O_2_ solubility and *pO*_2_ is the pO_2_.

The simulation geometry is based on a 24-well plate geometry with three distinct layers from bottom to top: (1) the bottom layer, which was defined either as a 50-μm-thick O_2_-permeable PMP membrane or as a completely non-O_2_-permeable membrane under a zero O_2_ permeability condition, (2) an active O_2_ consumption layer consisting of spheroids and culture media, and (3) the top culture media layer.

For the active O_2_ consumption layer, OCRs and related simulation parameters were defined on the basis of experimentally measured spheroid OCRs and measured O_2_ concentrations, as described in the following section *OCR of hepatocyte spheroids*. The spatial extent of O_2_ consumption within each well was modeled using a simplified homogenized consumption approach, in which the total O_2_ consumption of the seeded spheroids was distributed uniformly over the area occupied by the spheroids. In the dispersed distribution condition, O_2_ consumption was assigned over a circular area corresponding to the full culture area of the 24-well plate. In the central clustering condition, O_2_ consumption was assigned over a smaller circular area corresponding to the experimentally observed clustered spheroid region. The diameters of these clustered regions in the 200-and 600-spheroid conditions were determined from representative spheroid distribution images presented in the Results section. A complete list of simulation parameters is provided in **Supplementary Table S1**.

### OCR of hepatocyte spheroids

The OCR of hepatocyte spheroids was measured for use in computational simulations of culture media pO_2_ in wells seeded with hepatocyte spheroids. HepG2 spheroids were first generated on microwell plates as described in *HepG2 spheroid formation and culture conditions*. After spheroid formation, the cultures were pre-incubated at 37°C for 4 hours prior to OCR measurement. OCR was measured using a Seahorse XFe analyzer (Seahorse Bioscience, North Billerica, MA, USA). Approximately 100 spheroids were transferred to each well of a Seahorse XFe islet capture plate containing RPMI 1640 medium supplemented with 3 mM glucose. Basal OCR measurements were recorded every 7.5 minutes over a total measurement period of 45 minutes, and the average OCR was calculated from these measurements. The exact number of spheroids per well was determined by image analysis (cellSens, Olympus), and OCR values were normalized to the total spheroid volume. Data were obtained from four independent replicates and used to calculate the mean OCR.

### Cell proliferation assays of single cells

Cell proliferation in K562 suspension cultures and HepG2 monolayer cultures was quantified using a water-soluble tetrazolium salt-8 (WST-8) assay (DOJINDO Laboratories, Kumamoto, Japan) on culture days 0, 1, 2, and 3. WST-8 reagent was added to each well and incubated for 30 minutes. Absorbance of the sample was measured at 450 nm and 650 nm using a microplate reader (Infinite 200 PRO M Plex, Tecan Group Ltd., Männedorf, Switzerland). Absorbance values were normalized to the day 0 measurement to express relative values as percentages.

### Spheroid growth assays

Spheroid growth during culture was assessed by measuring spheroid size on culture day 2. Two days after culture in both flat plate and microwell plate studies, approximately 30 spheroids per condition were sampled and stained with 0.48 µM fluorescein diacetate (FDA; Sigma-Aldrich) to obtain fluorescence images using a Thunder Imaging System (Leica Microsystems, Wetzlar, Germany). Spheroid area was quantified using imaging software (cellSens, EVIDENT, Tokyo, Japan), and the estimated spheroid diameter was calculated assuming a spherical geometry.

### Viability assessment of spheroids using image analysis

Spheroid viability was analyzed using live/dead staining combined with a semi-automated image analysis method, as previously reported.^45, 46^ Approximately 100 spheroids per group were stained by incubation in phosphate-buffered saline containing 0.48 µM FDA and 15 µM propidium iodide (PI; Sigma-Aldrich) for 5 minutes in the dark at room temperature. Following staining, spheroids were imaged using the Thunder Imaging System (Leica, Wetzlar, Germany). For each condition, samples containing 10 to 20 spheroids were prepared for imaging. The total FDA-positive area representing live cells and the PI-positive area representing dead cells within all spheroids were quantified to determine overall spheroid viability using imaging software (cellSens, EVIDENT, Tokyo, Japan). Eight experimental replicates were performed.

### HMGB1 release assay

Cell damage in spheroid cultures was assessed by measuring the release of high mobility group box 1 (HMGB1), a ubiquitous nuclear protein that serves as a non-cell-type-specific biomarker of necrotic cell injury and inflammation.^47, 48^ HMGB1 released into the culture media was quantified using an HMGB1 Detection Kit (Chondrex, Woodinville, WA, USA) according to the manufacturer’s instructions. Culture media were collected on culture days 0 and 2, and HMGB1 concentrations were measured. Two to three independent experiments were performed.

### Gene expression assays (qPCR)

Total RNA was isolated from approximately 200 HepG2 spheroids collected from in vitro cultures at pre-culture (day 0) and post-culture (day 2) time points using TRI Reagent (Molecular Research Center, Cincinnati, OH, USA) in combination with the Direct-zol RNA Microprep kit (Zymo Research, Irvine, CA, USA) according to the manufacturers’ instructions. Extracted RNA was used for quantitative real-time PCR (qPCR) analysis following reverse transcription to generate cDNA. qPCR reactions were performed using TaqMan chemistry with gene-specific primer-probe sets obtained from Thermo Fisher Scientific (Waltham, MA, USA). Each reaction contained Quanta PerfeCTa qPCR Fastmix (95077-05K) (Beverly, MA, USA), TaqMan primer-probe mix, RNase-free water, and cDNA template in a final reaction volume of 10 µL per well. Reactions were assembled in 384-well plates and performed using a ViiA 7 Real-Time PCR System (Applied Biosystems, Thermo Fisher Scientific, Waltham, MA, USA). Briefly, the amplification steps involved: an initial enzyme activation and cDNA denaturation step (2 min at 50°C, then 2 min at 95°C) followed by 45 cycles of cDNA denaturation (15 s at 95°C) and probe annealing, elongation, and recording (1 min at 60°C). Technical triplicate reactions were performed for each sample for each probe. For each sample, the relative gene expression levels of hypoxia-associated genes (DNA damage inducible transcript 4 (*DDIT4*), egl-9 family hypoxia inducible factor 1 (*EGLN1*), solute carrier family 2 member 1 (*SLC2A1*), hexokinase 1 (*HK1*), and vascular endothelial growth factor A (*VEGFA*)) were normalized to the endogenous housekeeping gene *beta actin* (*ACTB*) using comparative analysis (= POWER(2,[Average Ct of *ACTB*] - [Ct value of technical replicate]) * 1000). Gene names, Gene IDs, and corresponding TaqMan assay IDs are listed in **Supplementary Table S2.**

## Statistical analysis

All statistical analyses were performed using JMP version 18.2.1 (SAS Institute, Cary, NC, USA). Data are presented as mean ± standard error of the mean (SEM), as indicated in the corresponding figure legends. Comparisons between two groups were conducted using unpaired two-tailed Student’s t-tests. For experiments involving three or more groups, one-way analysis of variance (ANOVA) was used, followed by the Tukey–Kramer honestly significant difference (HSD) post hoc test for multiple comparisons. Statistical significance was defined as P < 0.05.

For time-course dissolved O_2_ measurements, repeated measurements were analyzed using a linear mixed-effects model with experimental condition, time, and their interaction included as fixed effects. Repeated measurements obtained within each experimental run were modeled using an autoregressive covariance structure of order 1 (AR(1)) to account for temporal correlation. For cell proliferation assays, longitudinal WST-8 absorbance data were obtained and normalized to the day 0 measurement, which was defined as 100% based on the average absorbance value at day 0. The normalized data were analyzed using linear mixed-effects models. Culture day, O_2_ permeability condition (i.e., O_2_-permeable plate or non-O_2_-permeable plate), media replacement condition, and their interactions were included as fixed effects, as appropriate for each experiment. Repeated measurements obtained within each experimental run were modeled using an AR(1) covariance structure. Fixed-effect test results were reported in the corresponding figures and supplementary tables where applicable. For endpoint analyses, including equilibrated pericellular O_2_ tension, spheroid diameter, spheroid viability, HMGB1 release, and hypoxia-associated gene expression, Student’s t-tests or one-way ANOVA followed by Tukey–Kramer HSD post hoc testing were used, as appropriate for the number of groups compared. For qPCR experiments, technical triplicates were averaged before statistical analysis, and the averaged value from each independent experiment was treated as one biological replicate.

## Results

### Characterization of the O_2_-permeable plates

We quantified the O_2_ permeability of PMP in a flat membrane configuration (50 µm-thick), as summarized in **Table 1**. To generate a non-O_2_-permeable control while maintaining identical well geometry, a 100 µm PET gas barrier film was attached to the bottom of the PMP membrane. To support 3D spheroid culture, we fabricated microwell-structured PMP membranes (total thickness 135 µm, with a base thickness of approximately 50 µm at each microwell bottom; structural details are shown later). For reference, O_2_ permeability was also measured for conventional polystyrene culture plate material and PDMS.

**Table 1.**
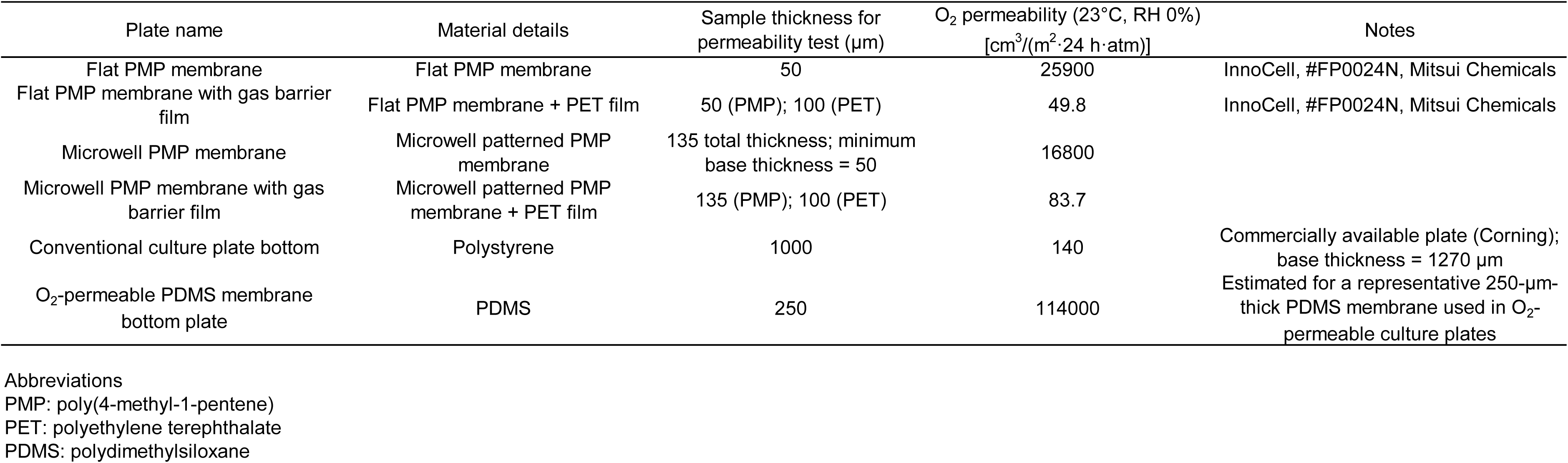
Oxygen permeability of the materials.

| Plate name | Material details | Sample thickness for permeability test (μm) | O<sub>2</sub> permeability (23°C, RH 0%)<br>[cm<sup>3</sup>/(m<sup>2</sup>·24 h·atm)] | Notes |
| --- | --- | --- | --- | --- |
| Flat PMP membrane | Flat PMP membrane | 50 | 25900 | InnoCell, #FP0024N, Mitsui Chemicals |
| Flat PMP membrane with gas barrier film | Flat PMP membrane + PET film | 50 (PMP); 100 (PET) | 49.8 | InnoCell, #FP0024N, Mitsui Chemicals |
| Microwell PMP membrane | Microwell patterned PMP membrane | 135 total thickness; minimum base thickness = 50 | 16800 |  |
| Microwell PMP membrane with gas barrier film | Microwell patterned PMP membrane + PET film | 135 (PMP); 100 (PET) | 83.7 |  |
| Conventional culture plate bottom | Polystyrene | 1000 | 140 | Commercially available plate (Corning); base thickness = 1270 μm |
| O<sub>2</sub>-permeable PDMS membrane bottom plate | PDMS | 250 | 114000 | Estimated for a representative 250-μm-thick PDMS membrane used in O<sub>2</sub>-permeable culture plates |

Abbreviations PMP: poly(4-methyl-1-pentene) PET: polyethylene terephthalate PDMS: polydimethylsiloxane

Overall, both the flat PMP membrane and the microwell-structured PMP membrane demonstrated substantially higher O_2_ permeability than conventional polystyrene culture plates, showing approximately 100 to 200-fold greater O_2_ permeability. In contrast, the PET gas barrier film effectively suppressed O_2_ diffusion, resulting in permeability values lower than those of conventional polystyrene culture plates. This design enabled direct comparison between O_2_-permeable and non-O_2_-permeable wells while preserving identical culture surface geometry for cells and spheroids.

### Flat O_2_-permeable bottom plates support high-density K562 suspension culture

We cultured K562 cells, a human chronic myelogenous leukemia suspension cell line, at high seeding density on flat O_2_-permeable bottom plates. **Figure 1A** summarizes the experimental design, including the culture duration and the comparison between O_2_-permeable and non-O_2_-permeable conditions.

**Figure 1.**
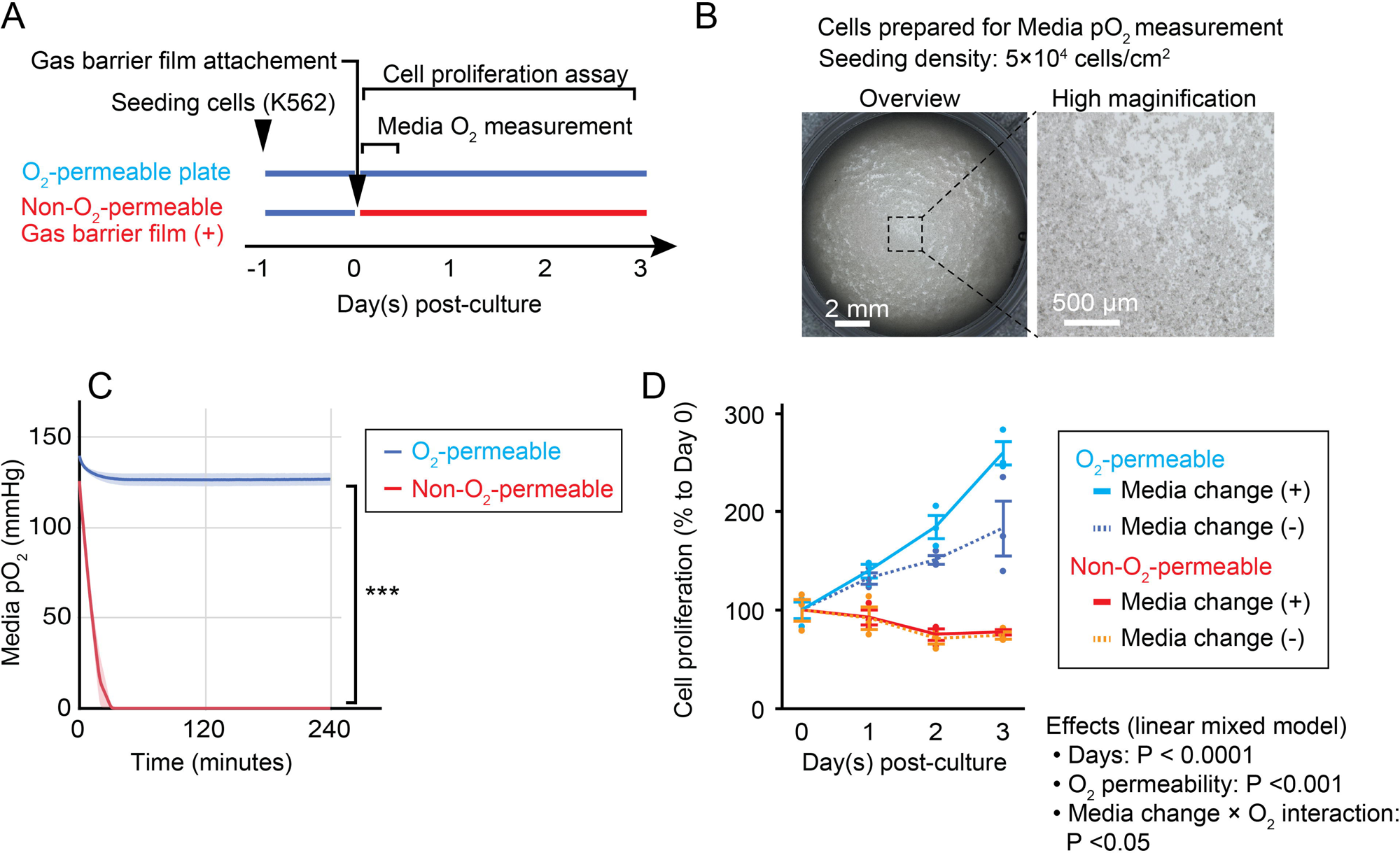
Oxygen (O_2_)-permeable bottom plates support high density suspension culture. (A) Experimental workflow for evaluating O_2_ availability and cell proliferation in O_2_-permeable and non-O_2_-permeable culture conditions. A human leukemia cell line, K562 cells, were seeded on flat O_2_-permeable plates, and gas barrier film was attached to the plate bottom to generate non-O_2_-permeable conditions. Media O_2_ measurements and cell proliferation assays were performed during the indicated culture period. (B) Representative images of K562 cells seeded for O_2_ measurement. Cells were seeded at a density of 5 × 10^5^ cells/cm^2^ one day prior to O_2_ measurement. Brightfield images show the distribution of high-density suspension cells within the well. Scale bars, 2 mm for overview images and 500 µm for magnified images. (C) Time-course measurement of media O_2_ tension following gas barrier film attachment. Dissolved O_2_ levels in the culture media were measured at one-minute intervals for 4 hours using a fluorescence-based O_2_ probe. Three independent measurements were performed in each condition. Data are presented as mean ± standard error of the mean (SEM) with shaded bands. Statistical comparisons were performed using a linear mixed-effects model. ***P < 0.0001. (D) Quantification of cell proliferation under O_2_-permeable and non-O_2_-permeable conditions with or without daily media replacement over three days of culture. Cell proliferation data obtained by water-soluble tetrazolium salt-8 (WST-8) absorbance were normalized to day 0 values and monitored. Each data point represents an independent experiment. Data are presented as mean ± SEM. N = 3. Statistical significance was determined using a linear mixed-effects model. Effects of culture day, O_2_ permeability, and media replacement interaction are summarized in the figure.

We seeded cells at a density of 5 × 10^5^ cells/cm^2^ in 24-well flat O_2_-permeable bottom plates one day prior to initiating O_2_ measurements (**Figure 1B**). We then attached a gas barrier film to the bottom of selected wells to create the non-O_2_-permeable condition and measured the dissolved O_2_ levels over a 4 hour period (**Figure 1C**). In non-O_2_-permeable wells, O_2_ tension rapidly decreased and approached near 0 mmHg within approximately 30 minutes after gas barrier film attachment. In contrast, O_2_-permeable plates maintained media O_2_ levels above 120 mmHg throughout the measurement period. Two-way ANOVA revealed a significant interaction between plate condition and time (F = 65.25, P < 0.0001), indicating that O_2_ dynamics differed markedly between O_2_-permeable and non-O_2_-permeable plates. We next evaluated cell proliferation over three consecutive days. K562 cells were seeded at a density of 2 × 10^5^ cells/cm^2^ in 96-well flat O_2_-permeable bottom plates one day before the start of the cell proliferation experiments. We selected this seeding strategy because cell proliferation rapidly reaches a plateau when cultures are initiated at excessively high densities. We then compared O_2_-permeable and non-O_2_-permeable plates under two culture conditions: daily media replacement and no media change (**Figure 1D**). Two-way ANOVA revealed that O_2_ permeability exerted a significant strong effect on cell proliferation from day 1 onward (P < 0.001). Media replacement showed no significant effect at day 1 (P = 0.6964) and day 2 (P = 0.0854), but became significant by day 3 (P = 0.0439). Importantly, significant interactions between O_2_ permeability and media replacement were observed at day 2 (P = 0.0340) and day 3 (P = 0.0314), indicating that media replacement enhanced proliferation primarily under O_2_-permeable conditions. In contrast, media replacement produced minimal improvement in non-O_2_-permeable plates, suggesting that O_2_ availability represents the dominant limiting factor for high density K562 cell proliferation. Detailed statistical analyses of cell proliferation under different O_2_ permeability and media replacement conditions are provided in **Supplementary Table S3**.

### Flat O_2_-permeable bottom plates maintain high O_2_ availability in HepG2 monolayer culture

We cultured HepG2 cells, a human hepatocellular carcinoma adherent cell line, as a 2D monolayer on flat O_2_-permeable bottom plates. **Figure S1A** outlines the experimental design, including cell seeding, attachment of a gas barrier film to generate non-O_2_-permeable conditions, measurement of media O_2_ levels, and subsequent analysis of cell proliferation.

We seeded HepG2 cells at densities of 4 × 10^4^ cells/cm^2^ one day before starting the experiment. We then measured dissolved O_2_ levels in the culture media immediately after attaching the gas barrier film (**Figure S1B**). In O_2_-permeable plates, media O_2_ levels remained stable at approximately 130–140 mmHg under both seeding density conditions. In contrast, non-O_2_-permeable wells showed a marked decline in O_2_ tension in the presence of cultured HepG2 cells, reaching progressively lower levels over time in a seeding density-dependent manner. On day 1, pericellular media O_2_ tension remained significantly higher in O_2_-permeable plates than in non-O_2_-permeable plates (**Figure S1C**; 133 vs. 35 mmHg, P < 0.0001). As a control, we also measured media O_2_ levels in the absence of cells and observed no difference between O_2_-permeable and non-O_2_-permeable plates (137 vs. 134 mmHg, P = 0.8007), confirming that cellular O_2_ consumption caused the O_2_ depletion observed in non-O_2_-permeable wells.

Next, we evaluated cell proliferation over three consecutive days under different O_2_ and nutrient conditions (**Figure S1D**). Representative photographs of the cultured cells demonstrated active proliferation during the early culture period, with cells approaching confluence within 2 days. Quantitative analysis further confirmed this pattern, with cell proliferation reaching a plateau or declining at later time points. In the no-media-change groups, proliferation declined after day 1, whereas media replacement sustained proliferation through day 2. Interestingly, O_2_ permeability did not significantly affect cell proliferation (**Figure S1E**). Linear mixed-effects analysis revealed media replacement significantly increased proliferation at both densities (P = 0.0303). In contrast, O_2_ permeability alone did not significantly affect proliferation (P = 0.4534).

Collectively, these results show that O_2_-permeable plates effectively maintain high O_2_ availability in 2D HepG2 cultures. In contrast, non-O_2_-permeable plates exhibited reduced pericellular O_2_ tension that stabilized at approximately 30 mmHg after equilibration during the first day of culture. Despite this O_2_ reduction, HepG2 cells maintained proliferation, indicating that O_2_ levels around 30 mmHg is sufficient to support 2D HepG2 cell growth. Under these conditions, nutrient replenishment exerted a stronger influence on proliferation compared to O_2_ availability.

### O_2_-permeable bottom plates improve spheroid growth in HepG2 spheroid culture

We next evaluated the performance of flat O_2_-permeable InnoCell^™^ plates for 3D HepG2 spheroid culture. Spheroid culture better recapitulates in vivo architecture but often experiences O_2_ limitation due to diffusion constraints, particularly within the spheroid core where severe hypoxia and necrosis can occur. Because spheroids contain densely packed cells, O_2_ demand increases substantially compared with 2D monolayer culture, which can also reduce O_2_ availability in the surrounding media. We therefore examined whether O_2_-permeable plates improve O_2_ supply and support spheroid growth under these conditions.

Unlike 2D monolayer cultures, spheroids can move within the well. As a result, media convection and minor vibrations in the incubator can cause spheroids to accumulate toward the center of the well, creating non-uniform O_2_ consumption and localized O_2_ gradients. To account for these effects, we evaluated two additional parameters in addition to O_2_ permeability: spheroid density (200 vs. 600 spheroids per well) and spatial distribution (dispersed vs. centrally clustered). **Figure 2A** summarizes the experimental workflow. HepG2 spheroids were first generated using a microwell platform and then transferred onto flat O_2_-permeable or non-O_2_-permeable plates created by attaching a gas barrier film. We then performed media O_2_ measurements and assessed spheroid growth and viability under these conditions. **Figure 2B** demonstrates that the microwell system produced uniformly sized spheroids within individual microwells during formation. FDA fluorescence imaging confirmed the formation of compact 3D spheroid structures before transfer to the flat plates. Macroscopic observation confirmed distinct spheroid distribution patterns between dispersed and centrally clustered conditions within individual wells of a 24-well plate at both low (200 spheroids per well) and high (600 spheroids per well) spheroid densities (**Figure 2C and 2D**).

**Figure 2.**
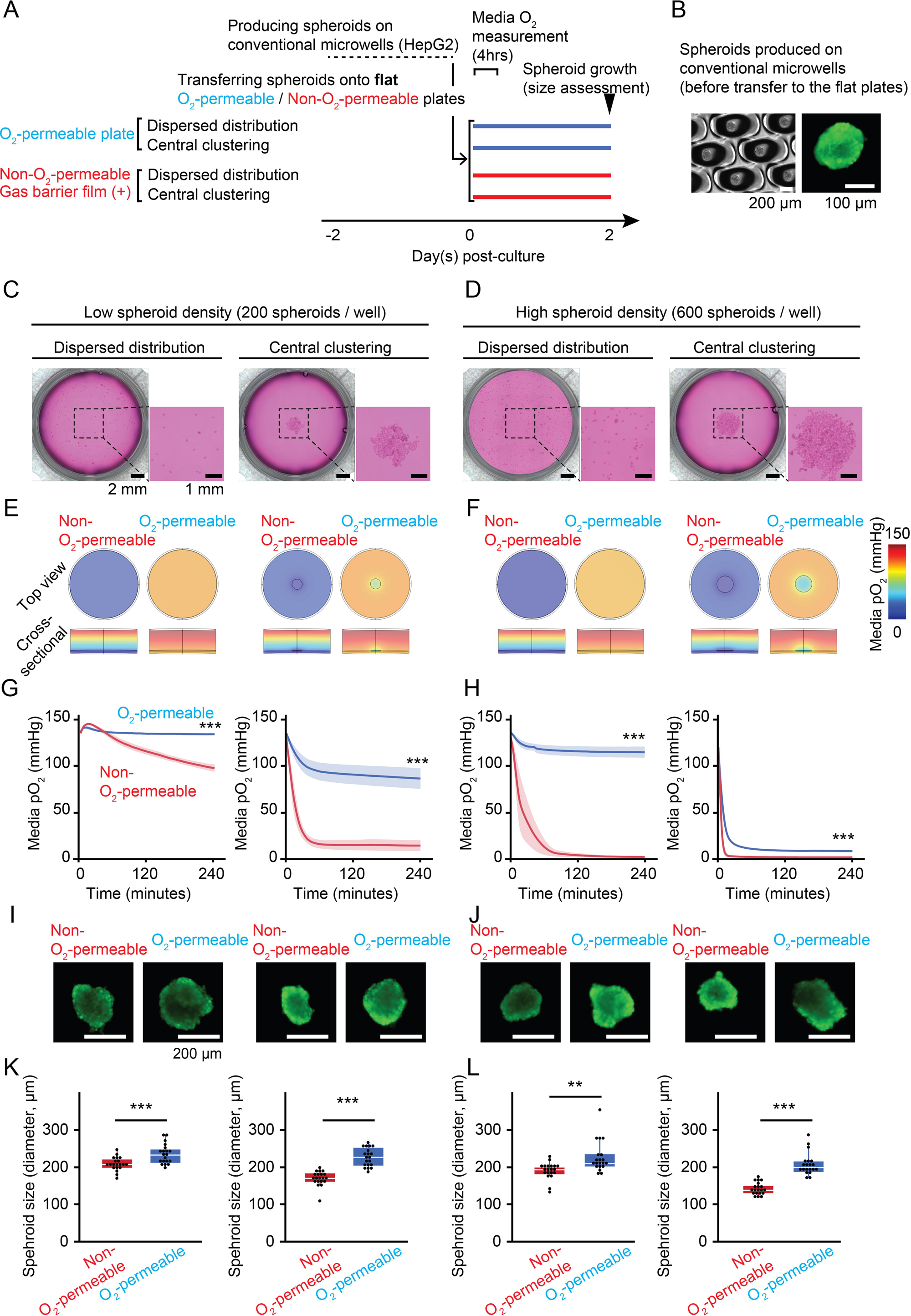
O_2_-permeable bottom plates improve spheroid growth in HepG2 spheroid culture. (A) Schematic illustration of the experimental workflow. HepG2 spheroids were first generated using a conventional microwell platform and subsequently transferred onto flat O_2_-permeable plates or non-O_2_-permeable plates created by attaching a gas barrier film to the plate bottom. Two spheroid densities (200 or 600 spheroids per well) and two spatial distributions (dispersed distribution or central clustering) were examined. Media O_2_ levels and spheroid growth were evaluated during subsequent culture. (B) Representative brightfield and fluorescent images of HepG2 spheroids formed in microwells prior to transfer onto flat culture plates. Brightfield and fluorescence images illustrate spheroid morphology and size before experimental manipulation. (C, D) Macroscopic images of wells showing spheroid spatial distribution after transfer to flat plates at low spheroid density (200 spheroids per well) and high spheroid density (600 spheroids per well) in a 24-well plate. Two distribution conditions were generated: dispersed distribution and central clustering. Left panels show an overview of the well, and right panels show enlarged images of the central region. Scale bars, 2 mm for overview images and 1 mm for enlarged views. (E, F) O_2_ diffusion simulations illustrating predicted O_2_ distribution within the culture wells under the indicated conditions. Top-view and cross-sectional simulations depict horizontal and vertical O_2_ gradients for O_2_-permeable and non-O_2_-permeable plates at low (200 spheroids per well) and high (600 spheroids per well) spheroid densities. (G, H) Time-course measurements of media O_2_ tension following spheroid transfer. Dissolved O_2_ levels were measured at one-minute intervals for 4 hours using a fluorescence-based O_2_ probe inserted into the culture media. Four independent experiments were performed, and data are presented as mean ± SEM with shaded bands. Statistical comparisons were performed using a linear mixed-effects model. ***P < 0.0001. (I, J) Representative fluorescence images of individual spheroids after 2 days of culture under the indicated conditions. Scale bars, 200 µm. (K, L) Quantification of spheroid diameter after 2 days of culture. Each data point represents an individual spheroid (N = 20 spheroids per condition). Box plots show the median (center line), interquartile range (box), and whiskers indicating the data range. Statistical significance was determined using Student’s t-test. **P < 0.01; ***P < 0.001.

We analyzed O_2_ distribution within the culture environment using O_2_ diffusion simulations (**Figure 2E and 2F**; **Supplementary Figure S2** shows the HepG2 spheroid OCR used for the simulation). Top-view and cross-sectional simulations illustrate horizontal and vertical O_2_ gradients within the wells. O_2_-permeable plates maintained higher O_2_ levels than non-O_2_-permeable plates across all conditions. Wells seeded with 600 spheroids exhibited lower O_2_ levels than wells seeded with 200 spheroids. Under both density conditions, centrally clustered spheroids generated pronounced O_2_ gradients with O_2_ tension decreasing toward the center of the well.

Direct measurement of media O_2_ levels further demonstrated O_2_ depletion dependent on both spheroid density and spheroid distribution, consistent with the simulation results (**Figure 2G and 2H**). The central clustering of spheroids at high spheroid density (600 spheroids per well) reduced O_2_ levels on flat O_2_-permeable plates due to the extremely high localized O_2_ consumption. In contrast, when spheroids were dispersed, wells seeded with 600 spheroids maintained media O_2_ levels above 100 mmHg. Together, these results indicate that both O_2_ permeability of the culture plate and spatial distribution of spheroids are critical determinants of O_2_ availability in spheroid cultures. Note that statistical analysis demonstrated significant differences between O_2_-permeable and non-O_2_-permeable conditions in each panel (P < 0.0001, linear mixed model).

Representative fluorescence images of individual spheroids after 2 days of culture are shown in **Figure 2I and 2J**, and quantitative analysis of spheroid diameters is summarized in **Figure 2K and 2L**. Across all conditions, spheroids cultured on O_2_-permeable plates were larger compared with those cultured on non-O_2_-permeable plates. Additionally, spheroids cultured under dispersed distribution conditions exhibited larger sizes compared with those cultured under central clustering configurations. Also, spheroids cultured at the lower seeding density (200 spheroids per well) exhibited larger diameters than those cultured at the higher density (600 spheroids per well). Within each density and distribution condition, O_2_-permeable plates consistently supported larger spheroid size compared with non-O_2_-permeable plates, with statistically significant differences observed across all conditions. Overall, these results indicate that O_2_ availability strongly influences spheroid growth under 3D culture conditions. Unlike 2D monolayer culture, spheroid systems impose substantially higher O_2_ demand due to the large number of cells packed within each aggregate and the uneven spatial distribution of spheroids within the well. Under these conditions, O_2_ delivery through the O_2_-permeable culture surface plays a critical role in supporting spheroid growth. These findings indicate that dispersed spheroid distribution is an important determinant for maintaining a more homogeneous O_2_ environment for highly O_2_-consuming spheroids.

### Spheroid culture on O_2_-permeable microwell plates prevents hypoxia-mediated spheroid injury in 3D structures

We have demonstrated that O_2_-permeable bottom plates support robust oxygenation, particularly during high seeding density cultures including 3D spheroids. However, spheroids can redistribute within the well during culture, often leading to central clustering. This clustering creates heterogeneous O_2_ microenvironments that are not fully compensated by O_2_ delivery from the plate bottom. To address this issue, we developed O_2_-permeable microwell plates to spatially control spheroid positioning and minimize localized O_2_ depletion. The microwell architecture maintained individual spheroids in separated positions during culture, thereby promoting a uniform spatial distribution within each well.

To target spheroid sizes commonly used for drug screening, cell therapy, and organoid applications (approximately 150 to 250 µm in diameter), we fabricated O_2_-permeable microwell plates with well dimensions of 350 µm in diameter and 85 µm in depth (**Figure 3A**). The membrane thickness at the bottom of each microwell was approximately 50 µm, allowing efficient O_2_ transport from the bottom of the plate directly into the culture media and spheroids (**Table 1**). We then applied the same spheroid culture workflow used in the flat plate experiments (**Figure 3B**). Importantly, the microwell system maintained spatial separation of spheroids even at high seeding density (600 spheroids per well in a 24-well plate) (**Figure 3C**) and prevented aggregation during culture. This configuration enabled a more uniform distribution of O_2_ consumption across the culture area and reduced the formation of localized hypoxic regions. Direct measurement of media O_2_ levels demonstrated sustained O_2_ availability in O_2_-permeable microwell plates, whereas O_2_ levels rapidly declined under non-O_2_-permeable conditions (**Figure 3D**; P < 0.0001, linear mixed model). Spheroid viability assessed by live/dead staining at day 0 and day 2 showed predominantly viable cells in spheroids cultured on O_2_-permeable microwell plates. In contrast, spheroids cultured on non-O_2_-permeable microwell plates with gas barrier film exhibited increased cell death, particularly in the spheroid core, consistent with hypoxia-mediated central necrosis (**Figure 3E**). Quantitative analysis confirmed significantly higher spheroid viability in O_2_-permeable microwell plates compared with non-O_2_-permeable controls at day 2 (P < 0.0001), and viability in O_2_-permeable conditions remained comparable to day 0 (P = 0.3724) (**Figure 3F**). We next assessed spheroid growth by measuring spheroid diameter as an indicator of cellular proliferation. Spheroids exhibited a significant increase in size from day 0 to day 2; importantly, spheroids in O_2_-permeable microwell plates showed a greater increase in diameter than spheroids cultured on non-O_2_-permeable microwell plates (P < 0.0001), indicating enhanced cell proliferation under improved O_2_ availability (**Figure 3G**). Consistent with the viability results, spheroids cultured on non-O_2_-permeable microwell plates released significantly higher levels of the cell damage and necrosis marker HMGB1 into the culture media after two days of culture (**Figure 3H**). In contrast, spheroids maintained on O_2_-permeable microwell plates released significantly lower HMGB1 levels, comparable to day 0 baseline values. Hypoxia-associated gene expression further supported the improved oxygenation provided by the O_2_-permeable microwell platform in transcriptional response level (**Figure 3I**). Spheroids cultured on non-O_2_-permeable microwell plates showed marked upregulation of multiple hypoxia-responsive genes, including *DDIT4*, *EGLN1*, *SLC2A1*, *HK1*, and *VEGFA*,^49–53^ after two days of culture. In contrast, spheroids maintained on O_2_-permeable microwell plates exhibited significantly lower expression of these genes, approaching levels observed at day 0. Collectively, these results show that O_2_-permeable microwell plate platform effectively prevents hypoxia-mediated spheroid injury and support sustained growth in high density cultures.

**Figure 3.**
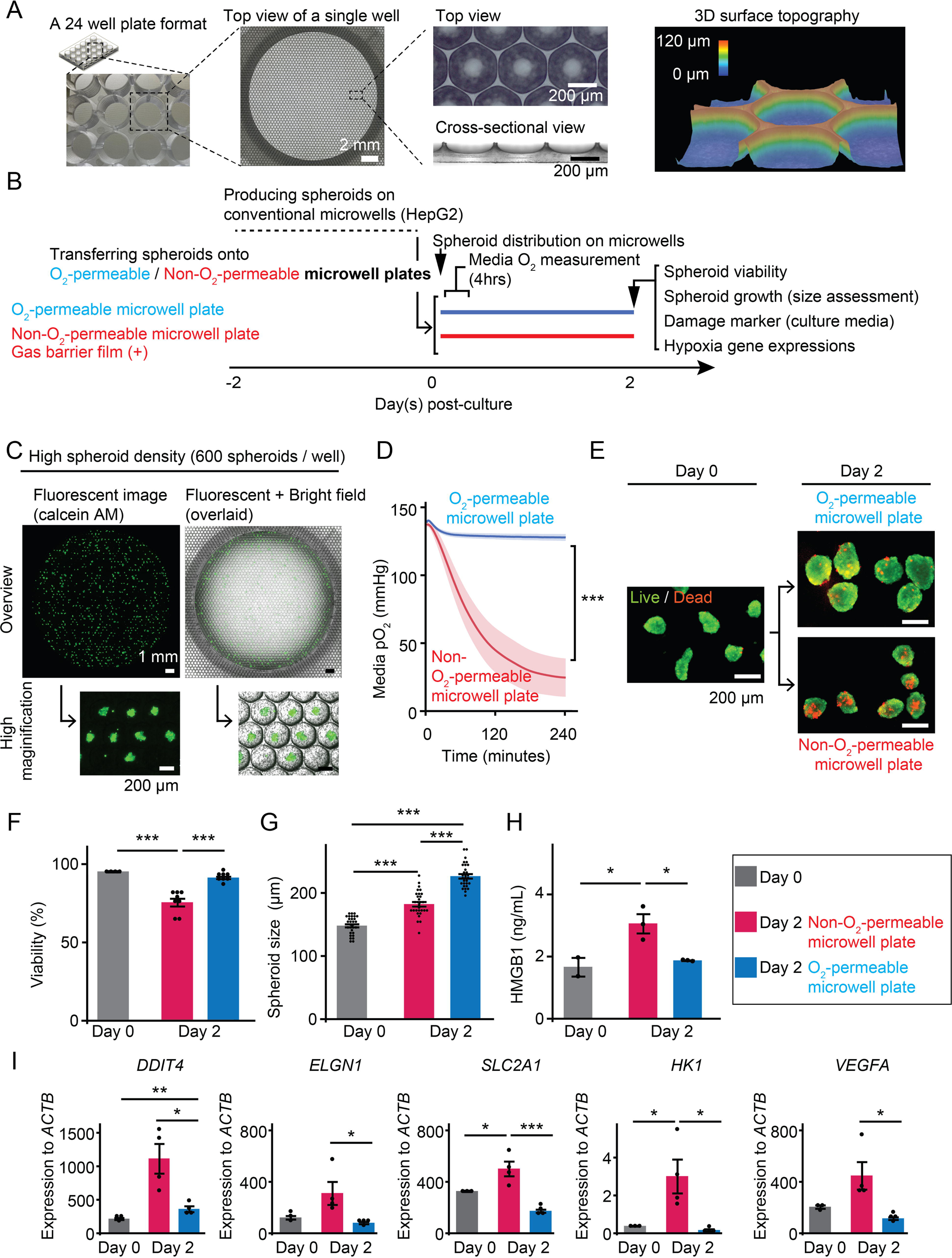
O_2_-permeable microwell plates prevent hypoxia-mediated injury in spheroid culture. (A) Structural characterization of the O_2_-permeable microwell plate. A 24-well plate format containing O_2_-permeable microwells is shown, including enlarged top-view images and cross-sectional views. Three-dimensional surface topography of the microwell structure was also obtained. Scale bars, 2 mm for the well overview and 200 µm for magnified images. (B) Experimental workflow for spheroid culture using the O_2_-permeable microwell platform. HepG2 spheroids were first generated using a conventional microwell system and subsequently transferred onto O_2_-permeable microwell plates or non-O_2_-permeable microwell plates generated by attaching a gas barrier film to the plate bottom. Subsequent analyses included media O_2_ levels, spheroid viability, spheroid growth, release of cell damage markers, and hypoxia-associated gene expression obtained from the cultured spheroids. (C) Representative fluorescence and brightfield images of spheroids cultured on O_2_-permeable microwell plates at high spheroid density (600 spheroids per well in a 24-well plate). Overview images and higher magnification views illustrate the spatial distribution of spheroids within the microwell array. Scale bars, 1 mm for overview images and 200 µm for magnified images. (D) Time-course measurement of media O_2_ tension following spheroid transfer to O_2_-permeable or non-O_2_-permeable microwell plates. Dissolved O_2_ levels were measured at one-minute intervals for 4 hours using a fluorescence-based O_2_ probe inserted into the culture media. Data are presented as mean ± SEM with shaded bands (N = 4 independent experiments). Statistical comparisons were performed using a linear mixed-effects model. ***P < 0.0001. (E) Representative live/dead fluorescence images of spheroids at day 0 and after 2 days of culture under the indicated conditions. Live cells are shown in green (fluorescein diacetate, FDA) and dead cells in red (propidium iodide, PI). Scale bars, 200 µm. (F) Quantification of spheroid viability at days 0 and 2 based on live/dead staining analysis. Each data point represents the assessment of the overall viability of approximately 20 spheroids. N = 8 per condition. Bars represent mean ± SEM. One-way ANOVA followed by the post-hoc Tukey–Kramer HSD test; ***P < 0.0001 by the post-hoc test analysis. (G) Quantification of spheroid diameter after 2 days of culture. Each data point represents an individual spheroid. N = 30 spheroids per condition. Bars represent mean ± SEM. One-way ANOVA followed by the post-hoc Tukey–Kramer HSD test; ***P < 0.0001 by the post-hoc test analysis. (H) Measurement of high mobility group box 1 (HMGB1) levels in the culture media after 0 and 2 days of culture. HMGB1 concentration was determined using ELISA. Bars represent mean ± SEM. Data were obtained from N = 3 independent experiments. ANOVA followed by the post-hoc Tukey–Kramer HSD test; *P < 0.05 by the post-hoc test analysis. (I) Quantitative real time PCR analysis of hypoxia-associated gene expression in spheroids after 2 days of culture under the indicated conditions. Gene expression levels of DNA damage inducible transcript 4 (*DDIT4*), egl-9 family hypoxia inducible factor 1 (*EGLN1*), solute carrier family 2 member 1 (*SLC2A1*), hexokinase 1 (*HK1*), and vascular endothelial growth factor A (*VEGFA*) were normalized to beta actin (*ACTB*). Bars represent mean ± SEM. Data were obtained from independent experiments repeated N = 4. Statistical significance was determined using ANOVA followed by the post-hoc Tukey–Kramer HSD test; *P < 0.05, **P < 0.01, ***P < 0.001 by the post-hoc test.

## Discussion

This study demonstrates the potential of PMP as a material for O_2_-permeable cell culture platforms. The use of PMP significantly enhanced oxygen availability within the culture system under high cell seeding density conditions. Notably, these effects were observed across both 2D and 3D culture systems, demonstrating the versatility of PMP-based platforms for a wide range of in vitro applications for O_2_-demanding cells. Importantly, even highly O_2_-demanding spheroids, characterized by densely packed cells within each aggregate, were effectively supported by the O_2_-permeable microwell platform. This was achieved through two complementary mechanisms: spatial control of spheroid distribution, which prevented spheroid aggregation and thereby limited localized regions of extremely high O_2_ consumption, and direct O_2_ supply through the bottom of the culture plate. By addressing both spheroid distribution and oxygen delivery, the platform enabled stable high-density spheroid culture. This O_2_-permeable microwell platform has substantial potential for high-throughput spheroid culture, accommodating approximately 8,000 microwells within a 35-mm-diameter culture area. Unlike bioreactor systems, which may introduce shear stress and require more complex operation,^54, 55^ this platform supports sufficient O_2_ delivery under static culture conditions without introducing bioreactor-associated shear stress.

Historically, a wide range of base materials has been used for cell-culture vessels. Polystyrene is currently the most commonly employed substrate due to its low cost, optical transparency, minimal drug adsorption, excellent amenability to surface modification, and high manufacturability. However, its inherently low O_2_ permeability imposes limitations in applications such as high seeding density culture, where O_2_ supply becomes a critical determinant of cell survival and function. Various O_2_-permeable materials have therefore been explored to enhance in vitro O_2_ delivery. Silicone elastomers, including PDMS, exhibit exceptionally high gas permeability and are easy to process, which has led to their use in cell culture platforms as membrane materials.^56–59^ Nevertheless, they present several limitations, including hydrophobic recovery that complicates stable surface modification, limited mechanical robustness, leaching of uncrosslinked oligomers, and substantial sorption of hydrophobic compounds.^32–35, 60, 61^ In particular, the absorption of hydrophobic molecules can compromise the reliability of drug-screening applications, raising concerns regarding their use in quantitative assays. Fluorinated ethylene propylene (FEP) is another gas-permeable material that is also used in cell-culture bags.^62^ Unlike PDMS, FEP exhibits very low molecular adsorption; however, fabrication of micro- or nano-scale surface structures remains technically challenging.^63, 64^ PMP has attracted increasing attention as an O_2_-permeable culture material because it combines high O_2_ permeability with optical transparency and mechanical strength and, importantly, exhibits lower drug sorption than PDMS.^36–41^ These properties have supported its use in culture vessels for drug evaluation using spheroid and tissue models.^44, 65, 66^

Several limitations of the present study should be acknowledged. First, although the major culture formats were covered, including suspension culture, 2D monolayer culture, and 3D spheroid culture, a limited number of cell types were covered. In addition, spheroid experiments were performed using pre-formed spheroids generated by a conventional platform rather than forming spheroids directly on the O_2_-permeable microwell plate. This approach simplified evaluation of the effect of O_2_ supply on spheroid maintenance; however, efficient O_2_ delivery may also influence spheroid formation from single cells seeded directly onto the microwell platform. Further validation across a broader range of cell types, particularly those with high metabolic and OCRs, will be necessary to establish the general applicability of the system.

Another important consideration is the optimization of microwell design, including diameter and depth. In this study, microwells with a diameter of 350 µm and a depth of 85 µm were used, supporting spheroids up to approximately 350 µm in diameter, with typical spheroid sizes of 150–200 µm. At the tested seeding density, approximately one spheroid was captured per microwell, and aggregation was not observed. However, higher seeding densities may increase the likelihood of multiple spheroids occupying a single microwell, indicating that optimization of seeding density and microwell dimensions may be required for specific applications. The selected depth facilitated efficient spheroid capture and recovery; however, deeper microwells may increase retention while complicating retrieval and promoting air bubble trapping. Therefore, the ratio of microwell diameter to depth represents an important design parameter that may require customization. In addition, the PMP material used in this platform is compatible with fabrication methods such as heat embossing, enabling flexible production of microwells with varied geometries. This adaptability provides opportunities for further optimization of the platform to accommodate diverse spheroid sizes, cell types, and experimental conditions.

In conclusion, the PMP-based O_2_-permeable microwell platform combines efficient O_2_ delivery with spatial control of spheroid distribution, supporting stable high-density 3D culture. This platform may provide a scalable and versatile approach for high-throughput applications, including drug screening, tissue engineering, and regenerative medicine.

## ACKNOWLEDGMENTS

None.

## AUTHOR CONTRIBUTIONS

J.T. and H.Komatsu conceived and designed the research; J.T., H.Kato, H.N.Z., K.I., S.M., and H.Komatsu performed experiments; J.T., H.Kato, H.N.Z., K.I., and H.Komatsu analyzed data; J.T., H.Kato, H.N.Z., H.T.K., K.I., S.M., K.E., and H.Komatsu interpreted results of experiments; J.T., K.I., and H.Komatsu prepared figures; J.T., H.N.Z., K.I., and H.Komatsu drafted the manuscript; H.Kato, H.T.K., K.I., S.M., K.E., and H.Komatsu edited and revised the manuscript; J.T., H.Kato, H.N.Z., H.T.K., K.I., S.M., K.E., and H.Komatsu approved the final version of the manuscript.

## STATEMENTS AND DECLARATIONS

### Ethical considerations

Ethical approval was not required because this study used established human cell lines obtained from established cell repositories and did not involve human participants or identifiable human participant data.

### Consent to participate

Not applicable because no human participants were enrolled in this study.

### Consent for publication

Not applicable because this study does not contain identifiable data, images, or information from individual human participants.

### Human rights

Not applicable because no human participants were involved in this study.

### Declaration of conflicting interest

Jun Takahashi, Kozo Iiba, Shunya Mizuno, and Katsuhiro Esashika are employees of Mitsui Chemicals, Inc. The University of California, San Francisco receives research support from Mitsui Chemicals, Inc. through a sponsored research agreement (principal investigator: Hirotake Komatsu; co-investigator: Hiroyuki Kato). The remaining authors declare no potential conflicts of interest.

## Funding

This work was supported in part by the Nora Eccles Treadwell Foundation (no grant number, to H. Komatsu), Breakthrough T1D Foundation (3-SRA-2025-1634-S-B and 1-SRA-2025-1635-S-B, to H. Komatsu), a postdoctoral fellowship from the California Institute for Regenerative Medicine (EDU4-12772, H.N.Z.), and a research grant from Mitsui Chemicals, Inc. (to H. Komatsu).

## Data availability

Data are available upon request.

## SUPPLEMENTAL MATERIAL

The following supplemental materials for this article are available online

Supplementary Figure S1

Supplementary Figure S2

Supplementary Table S1.

Supplementary Table S2

Supplementary Table S3

## Notes

### Summary of Updates

Corrected garbled text, Figures 2 and 3, author affiliation information, and the "Declaration of conflicting interests" statement. Added titles to Supplementary Tables S1 and S2.

