## Supplementary figures and images for "Oxygen-permeable microwell plates for high-throughput spheroid culture"

### Supplementary Figure S1

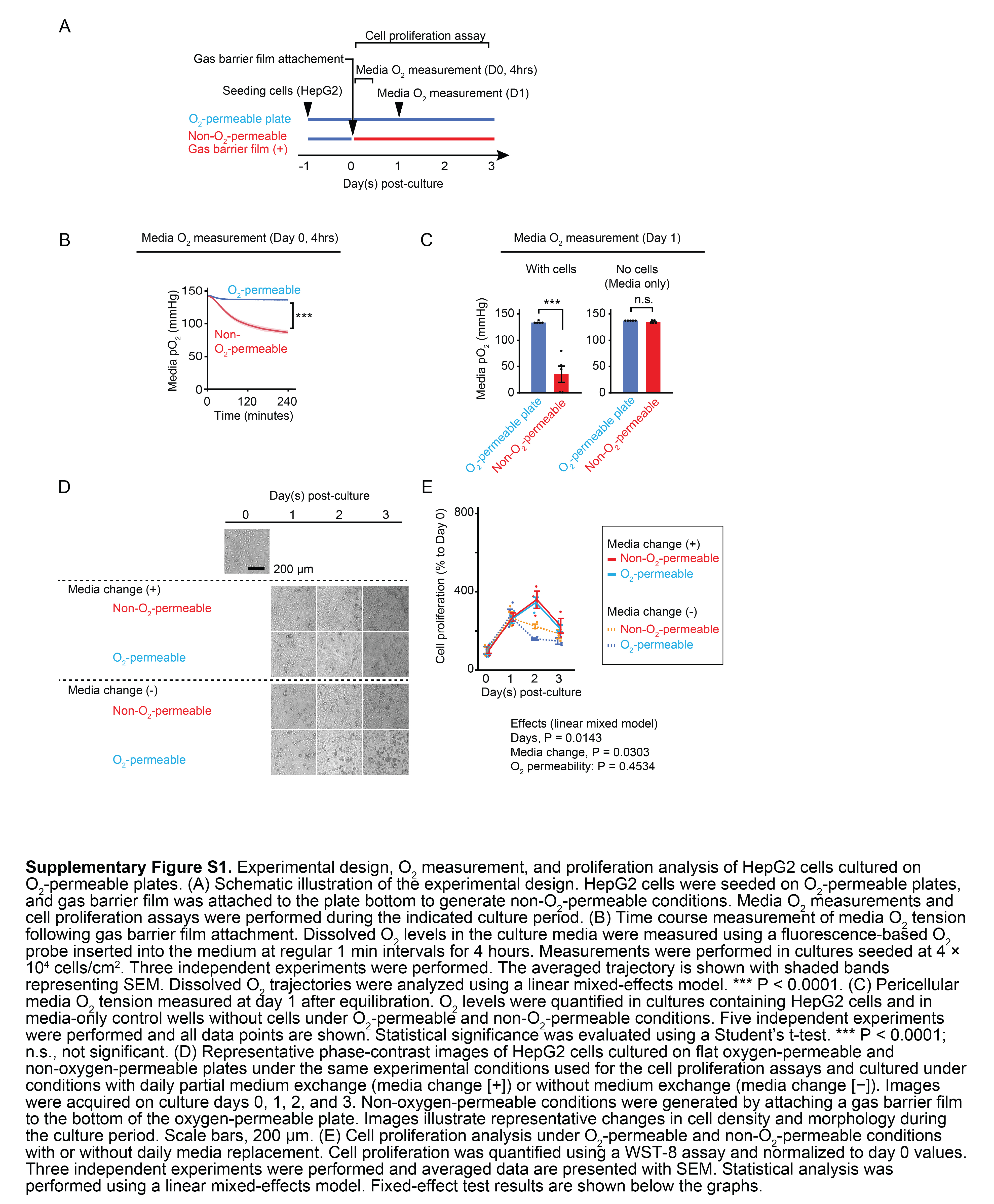

### Supplementary Figure S2

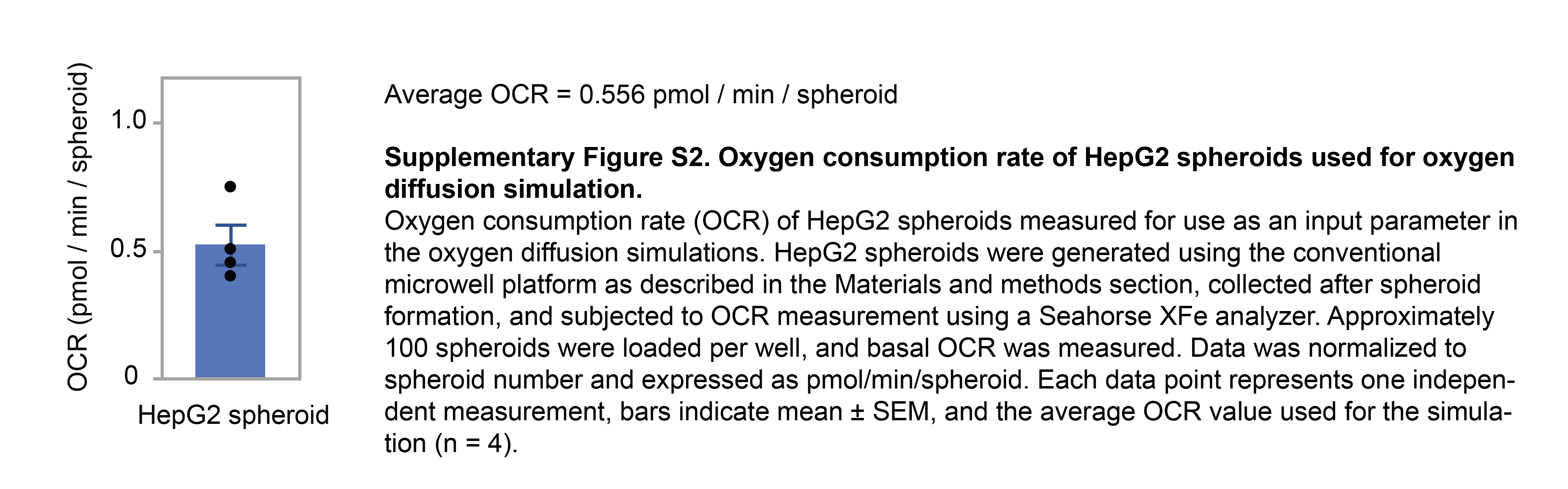
