## Supplementary Table S1 for "Oxygen-permeable microwell plates for high-throughput spheroid culture"

**Supplementary Table S1. A complete list of simulation parameters.**

| Description | Abbreviation | Value | References |
|---|---|---|---|
| Diffusivity of oxygen in medium | D_medium | 2.8×10^(-9) [m^2 ⟦·s⟧^(-1)] | [1] |
| Diffusivity of oxygen in poly(4-methyl-1-pentene) (PMP) | D_PMP | 1.6×10^(-9) [m^2 ⟦·s⟧^(-1)] | Measured |
| Solubility of oxygen in medium | S_medium | 9.7 ×10^(-6) [s^(2·) mol·kg^(-1) ⟦·m⟧^(-2)] | [2] |
| Solubility of oxygen in PMP | S_PMP | 7.6 ×10^(-6) [s^(2·) mol·kg^(-1) ⟦·m⟧^(-2)] | Measured |
| Michaelis oxygen constant of spheroid | K_m | 7.4 ×10^(-3) [mol·m^(-3)] | [3] |
| Maximum oxygen consumption rate of 200 spheroids | OCR_sph200_max | 6.5×10^(-5) [mol·m^(-3)·s^(-1)] | Measured |
| Maximum oxygen consumption rate of 600 spheroids | OCR_sph600_max | 1.9×10^(-4) [mol·m^(-3)·s^(-1)] | Measured |
| Atmospheric oxygen partial pressure | pO_2 | 151 [mmHg] | - |
| Diamter of 200 spheroid clusters in a culture dish | R_sph200 | 1167 [μm] | Measured |
| Diamter of 600 spheroid clusters in a culture dish | R_sph600 | 1889 [μm] | Measured |
| Temperature | T | 37 [℃] | [4] |
