## Supplementary Table S2 for "Oxygen-permeable microwell plates for high-throughput spheroid culture"

**Supplementary Table S2. Gene names, Gene IDs, and corresponding TaqMan assay ID.**

| Gene Symbol | Official Full Name | Gene ID | Taqman Assay ID |
|---|---|---|---|
| DDIT4 | DNA damage inducible transcript 4 | 54541 | Hs00430304_g1 |
| EGLN1 | egl-9 family hypoxia inducible factor 1 | 54583 | Hs00254392_m1 |
| SLC2A1 | solute carrier family 2 member 1 | 6513 | Hs00892681_m1 |
| HK | hexokinase 1 | 3098 | Hs00175976_m1 |
| VEGFA | vascular endothelial growth factor A | 7422 | Hs00900054_m1 |
| ACTB | actin beta | 60 | Hs01060665_g1 |
