## Supplementary Table S3 for "Oxygen-permeable microwell plates for high-throughput spheroid culture"

**Supplementary Table S3. Statistical analysis of cell proliferation under different oxygen permeability and media replacement conditions.**

| Day | Factor | p value |
|---|---|---|
| Day 1 | O2 permeability | 0.0007 |
| | Media change | 0.6964 |
| | Interaction (MC × O2 permeability) | 0.6277 |
| Day 2 | O2 permeability | <0.0001 |
| | Media change | 0.0854 |
| | Interaction (MC × O2 permeability) | 0.0342 |
| Day 3 | O2 permeability | <0.0001 |
| | Media change | 0.0439 |
| | Interaction (MC × O2 permeability) | 0.0314 |

Abbreviations: MC, media change.

Statistical test: Two-way ANOVA was performed separately for each culture day to evaluate the effects of oxygen permeability, media replacement, and their interaction.
